# Learning a reversed bicycle disrupts predictive control and induces interference with the normal bicycle

**DOI:** 10.64898/2026.05.08.723825

**Authors:** Pascal Nietschmann, David W. Franklin

## Abstract

Motor skills such as bicycle riding are considered robust and transferable across bicycle types. However, when the steering direction is inverted (reversed bicycle) control is disrupted to the extent that the bicycle cannot be ridden. With sufficient practice, the reversed bicycle can be learned, but this learning appears to produce impairment of normal bicycle riding suggesting modification of this long-established motor memory. Here we investigate the learning process of riding a reversed bicycle over four days of practice, while repeatedly assessing normal bicycle performance to measure any potential interference. Introduction of the reversed bicycle disrupted predictive control, reflected in a consistently increased time lag in the steering-roll coupling during reversed bicycle trials. This increase in delay suggests that predictive behavior in normal bicycle riding cannot be transferred to the reversed bicycle. With training, some participants successfully learned to ride the reversed bicycle by gradually reorganizing this coupling, whereas others failed to acquire this inverted coupling. Notably, even short-term exposure to the reversed bicycle interfered with normal bicycle riding, reducing distance ridden and increasing variability in steering rate. Together, we show that even a highly practiced whole-body motor skill is susceptible to rapid interference when control dynamics are altered.

## Introduction

Riding a bicycle is an extremely common motor skill that many humans perform, despite the complexity of the skill^1^. Once learned, such complex skills are retained robustly^2^, so that even after years without practice we can effortlessly start to ride a bicycle. We can generalize our learning from one task, (e.g. riding a bicycle) to similar tasks (e.g. motorcycle riding)^3–6^. Even when we change between bicycle types, we quickly adjust to such a change^7^. In contrast, if we change the direction of the steering by 180°, so that a rightward handlebar turn leads to a leftward wheel turn and vice versa, this reversed bicycle becomes initially unrideable. Studies have shown that some participants can successfully learn to ride this reversed bicycle after extensive practice^8–13^. Interestingly, during the learning process of this reversed bicycle, Magnard and colleagues found that there is substantial interference when switching back from the reversed bicycle to the normal bicycle, resulting in participants requiring several attempts to ride a normal bicycle again^13^. This interference effect is not commonly seen in many everyday activities, but has often been found in laboratory tasks where the visual feedback or dynamics are reversed or transformed for the same actions^14–19^. Here we further examine how interference arises in a very well learned real-world task (riding a normal bicycle) through the introduction of the reversed bicycle.

To quantify both how we learn the reversed bicycle as well as measuring the interference with normal bicycle riding, it is important to examine how a bicycle is balanced. Though the general design of bicycles has not changed significantly over the past 100 years^20^, it is still not fully understood how we control and balance a bicycle^1,21–25^. One major strategy to keep the bicycle upright is steering into the same direction that the bicycle tilts^26,27^. That is, when the bicycle tilts (bicycle roll), we steer into the same direction to cause a turn that generates a pseudo-centrifugal force redirecting the center of mass back above the base of support^28,29^. While steering remains the dominant strategy, lateral displacement of the upper body can also shift the center of mass, which influences bicycle roll in an indirect manner via the front frame^30^. Another mechanism to shift the center of mass is moving our contralateral knee outwards, which is particularly effective during riding with low velocities^27^. Indeed, the combination of trunk and knee movement, along with the self-stabilizing effect of the bicycle, allows us to ride a bicycle without the use of hands on the handlebar^31^.

There are many different parameters that are used to quantify riding and balance performance. The steering performance can be quantified by the variability in the steering angle or steering rate^32–36^. Similarly, lower variability of bicycle roll angle or rate, is a measure of roll stability and balance control^35–37^. Variability in trunk lean is also a measure of balance performance, as it reflects upper-body strategies to maintain stability^33,34^. In addition, overall riding performance can be assessed with the riding trajectory, where reduced lateral deviation reflects improved control of the bicycle^35^.

Cyclists constantly adjust steering to regulate bicycle roll and keep the bicycle upright. As a steering action follows a bicycle tilt, these two parameters are closely coupled (correlation coefficient 0.6-0.8) with a time lag of 90-140 ms^33,34,37–39^. The fact that this correlation is less than one indicates that other factors also contribute to balance such as trunk lean^1^. This short time delay between bicycle roll and steering suggests that precise, predictive control is necessary to maintain balance^23,25^. Such predictive control may be dramatically affected when the mapping between actions and outcomes is reversed, such as in the reversed bicycle. Reversing the mapping increases reaction times^40,41^ and produces initial corrections to the wrong direction, even after extended practice^40,42^. Given the limited time available for steering corrections with a bicycle before falling over, these findings help explain why the reversed bicycle is particularly challenging to initially ride.

While several studies have shown that the reversed bicycle is learned slowly^8–12^, only Magnard and colleagues showed that it also introduces interference with normal bicycle riding^13^. They showed that this interference increased from day 5 to day 9, and produced low frequency oscillations in the handlebar steering. However, it is unknown how quickly interference on the normal bicycle starts to appear. Does it already appear after initial training with the reversed bicycle? How does interference develop with further training of the reversed bicycle? Maintaining stability in bicycle riding depends not only on steering behavior, but also on adjustments in the bicycle roll, trunk movement, and riding trajectory. Together, these measures might provide evidence of interference and, more critically, could help quantifying the learning of the reversed bicycle. To address these issues, we probe the emergence and time course of interference twice per day with interleaved catch trials and a range of kinematic measures. In addition, we extend current approaches to quantify reversed bicycle learning by examining balance-related performance measures throughout the experiment and by investigating the reorganization of steering-roll coupling during adaptation to the reversed bicycle.

## Results

Twenty participants came to the laboratory for four days and completed the experiment designed to assess learning on a reversed bicycle and potential interference with normal bicycle riding. Bicycle and participant kinematics were assessed using a 3D motion capture system (Fig. 1a), with reflective markers placed on the bicycle and body segments (Fig. 1b). From these data, we computed the distance achieved, the steering angle, bicycle roll angle, trunk lean, and lateral knee movement. We quantified steering, balance, and riding performance as steering rate variability (standard deviation of the steering rate), roll rate variability, trunk lean rate variability, lateral knee velocity variability, and straight-line deviation (root mean square of the lateral deviation from a straight line). Furthermore, to assess the coupling between bicycle roll and steering, we calculated the cross correlation between these two parameters and investigated the peak correlation coefficient and its time lag. Due to the limited capture volume, forward riding distance was limited to six meters per trial to ensure comparability and successful marker tracking. On the first day, participants started with the normal bicycle and first completed five baseline trials. This was followed by five pre-tests on the reversed bicycle. These five pre-tests with the reversed bicycle were performed at the start of all following days of the experiment. After the pre-tests, training consisted of 20 minutes of free riding per day on the reversed bicycle. We interleaved single, normal bicycle trials (catch trials) to assess interference with normal bicycle riding throughout the experiment. At the end of each day, participants performed five post-tests with the reversed bicycle (Fig. 1c). Finally, on the fourth day, participants were reintroduced to the normal bicycle with a series of testing trials and training.

**Figure 1.**
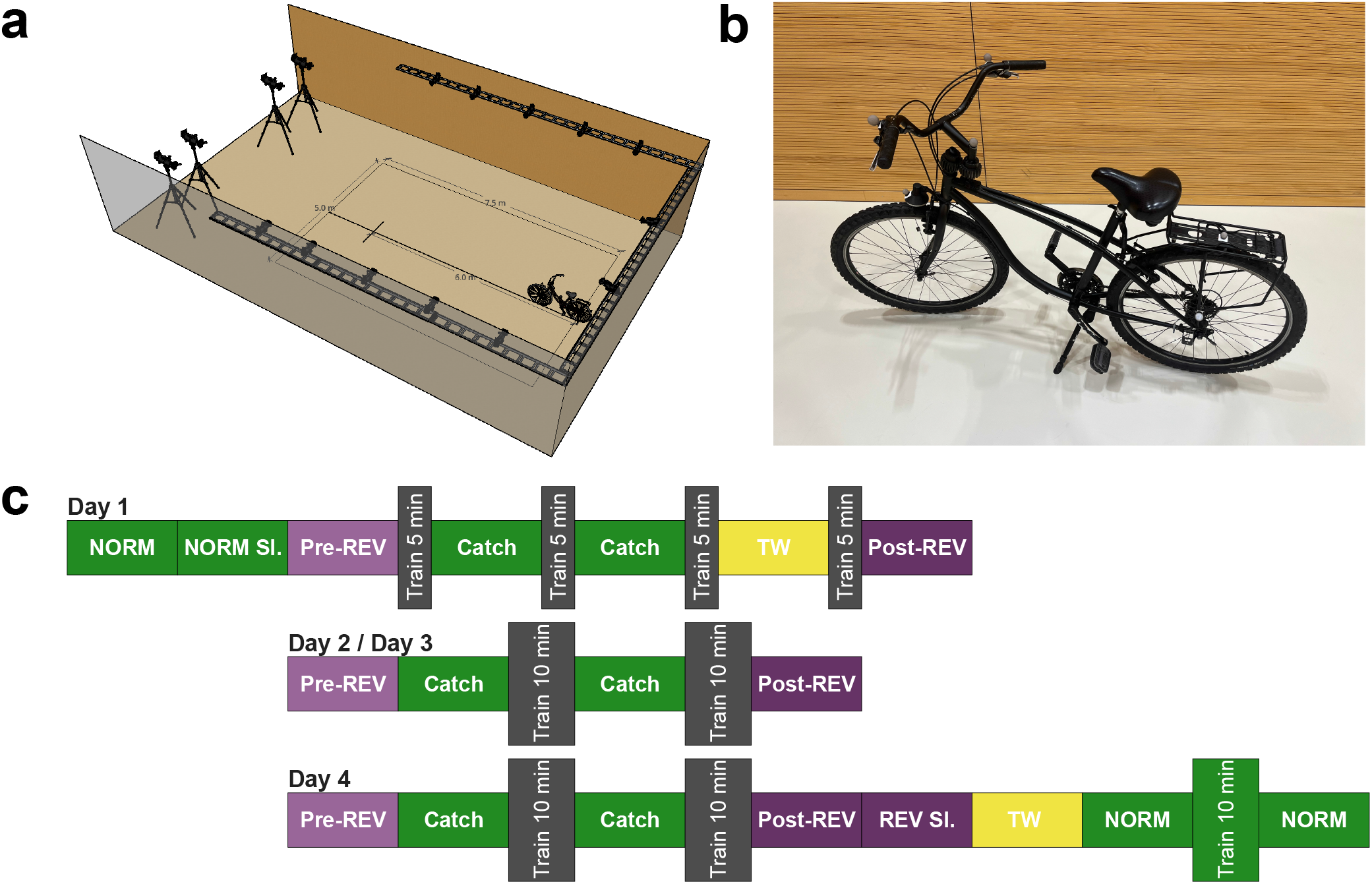
Experimental Design. (a) 3D motion capture setup in a sports hall with a capture volume: 7.5 × 5.0 meters. Six-meter line is indicated with a short black line. (b) The reversed bicycle with reflective markers. (c) Experimental protocol across four days: Baseline trials (tr) with the normal bicycle (NORM), pre- and post-tests with the reversed bicycle (Pre-REV, Post-REV), free training sessions (Train) with the reversed bicycle (gray) and with the normal bicycle (green), and reversed bicycle trials with training wheels (TW). Sl. indicates slalom. Five trials were performed in each block except in catch trials (1 trial) and in TW trials (1 trial).

### Distance-based performance

First, we examined how the introduction of the reversed bicycle affected the forward riding distance of the participants with the reversed bicycle, and whether it led to interference with the normal bicycle. Distance was assessed as either the full six meters in the case of a successful trial or the first point when participants placed their foot on the ground. During baseline testing with the normal bicycle, participants consistently achieved the full six meters. In the first five pre-test trials, when participants stepped on the reversed bicycle for the first time, the average distance achieved was less than one meter (0.90 ±0.07 m, Fig. 2a). This distance increased with training to a final level of 4.09 ±0.47 m in the post-tests on day four. An ANOVA yielded a significant effect of learning both within days (pre-post: F_1,19_=50.063, p<0.001,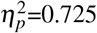) and across days (days: F_1.25,23.69_=35.198, p<0.001, 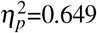). Analysis of individual results revealed substantial differences in the amount of distance achieved (Fig. 2b+c) ranging from 6.50 to 42.51 meters. We ranked and color coded participants based on the sum of distances from the pre- and post-tests (Fig. 2b) with darker colors indicating the participants with further distances ridden with the reversed bicycle.

**Figure 2.**
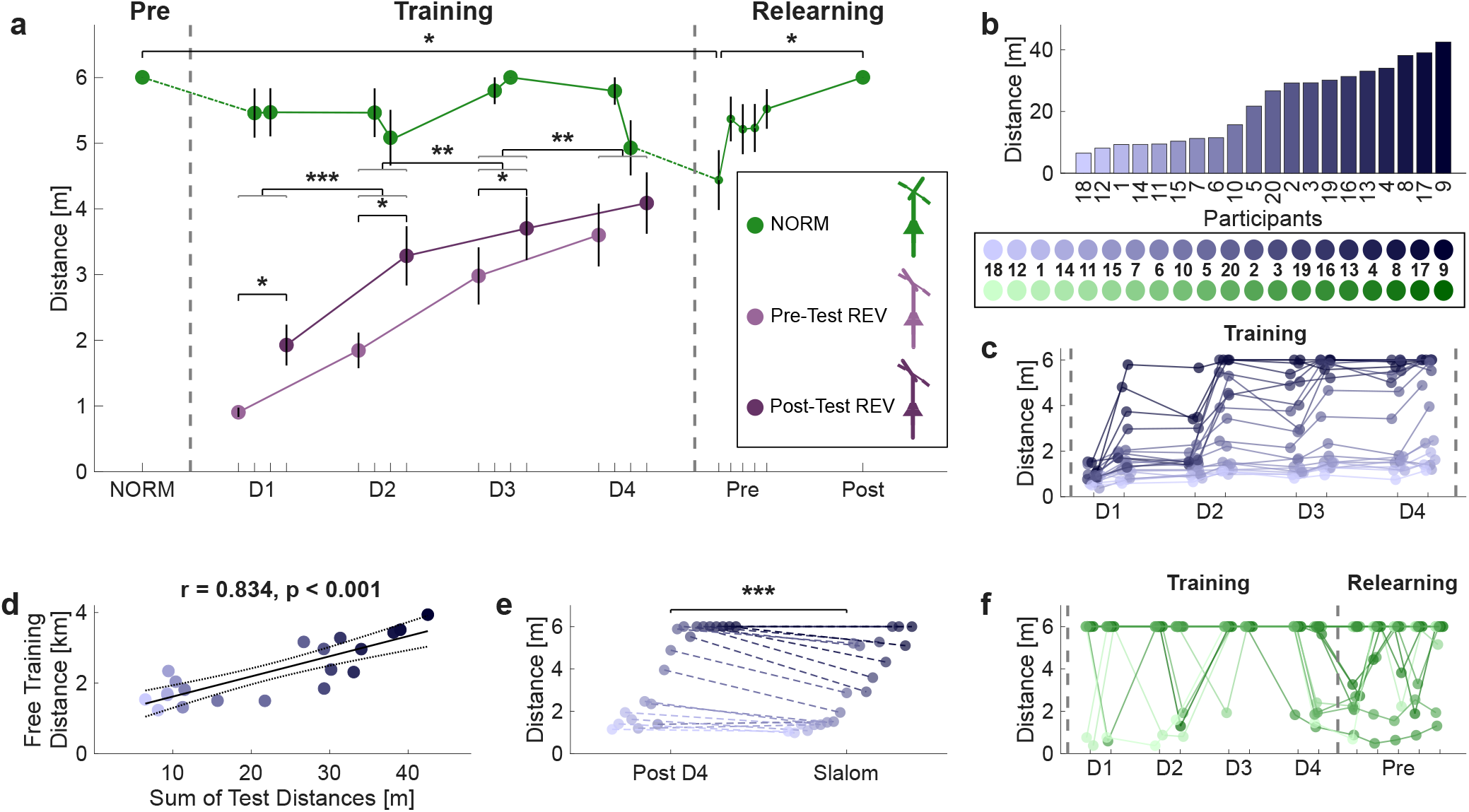
Forward distances ridden on the normal and reversed bicycle across the experiment. (a) Mean distance achieved across participants over the course of the experiment for normal bicycle trials (green), reversed bicycle pre-tests (light purple) and reversed bicycle post-tests (dark purple). Dashed lines distinguish experimental phases. Asterisks denote significant differences based on pairwise comparisons (* p<0.05; ** p<0.01; *** p<0.001). When multiple pairs are significant, a gray bracket is used to group them.(b) Summed distance achieved by each participant on the reversed bicycle across both the pre- and post-tests, sorted by total distance (Color gradient from light to dark blue). (c) Individual distances in reversed bicycle pre- and post-tests, with color gradient indicating individual participants (see legend).(d) Relationship between summed training distance in the free training sessions (20 minutes per day) and summed total test distance. Solid line shows linear fit, the dotted lines indicate the 95% confidence interval.(e) Comparison of final straight-line and slalom tests with the reversed bicycle. (f) Individual distances in catch trials and pre-tests of the relearning phase. Individuals are indicated by colors where darker green indicates participants that had larger total distance on the reversed bicycle (see legend in b).

To assess if participants who rode more distance during free training also achieved more distance in the pre- and post-tests, we correlated the sum of pre- and post-test distances to the total distance ridden in the free training sessions which revealed a strong positive linear correlation (r=0.834, p<0.001, Fig. 2d). We observed no significant correlations between questionnaire measures on physical activity level, bicycle riding frequency, or riding distance with the sum of test distances on the reversed bicycle (Supplementary Fig. S1).

To assess to what degree participants were able to actively control the reversed bicycle, we asked them to ride in a slalom on day four. Distance achieved in these slalom trials was less compared to straight-line riding on day four post-tests (Wilcoxon signed-rank, W=198, p<0.001, Fig. 2e), but with a strong correlation between these measures across participants (r= 0.911, p<0.001).

As the learning of the reversed bicycle could interfere with the normal bicycle riding, we examined the distance ridden on the normal bicycle throughout the experiment. While many participants were able to ride the full six meters on the normal bicycle during the catch trials, there were almost always some participants that stepped down earlier (Fig. 2f). An ANOVA examining the distance ridden on the normal bicycle yielded a significant main effect (F_1.510,28.686_=5.765, p=0.013, 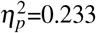). Post-hoc comparison revealed significantly less distance achieved in the first relearn trial at the end of day four compared to both the initial baseline trials (t_19_=3.434, p=0.017, Cohen’s d=1.187) and the final post relearn trials (t_19_=−3.434, p=0.017, Cohen’s d=−1.187).

Overall, the distance metric revealed two key findings. First, participants were gradually able to learn to ride the reversed bicycle, although performance varied substantially across participants. Second, as participants improved on the reversed bicycle, they were occasionally no longer able to ride 6 meters with the normal bicycle, suggesting interference with normal bicycle riding.

### Changes in steering control

While distance provides a rough overview of performance, we can also assess specific measures over the controllability of the bicycles. Control over the steering was assessed by investigating steering rate variability, calculated as the standard deviation of the steering rate. In baseline trials with the normal bicycle, the steering rate variability was 30.37 ±1.51 °/s. In both pre-tests and post-tests with the reversed bicycle the steering rate variability was higher, with no clear trend across days (Fig. 3a). An ANOVA on the steering rate variability with the reversed bicycle yielded no main effect within days (F_1,19_=1.740, p=0.203, 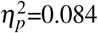) or across days (F_1.892,35.945_=0.608, p=0.541, 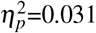).

**Figure 3.**
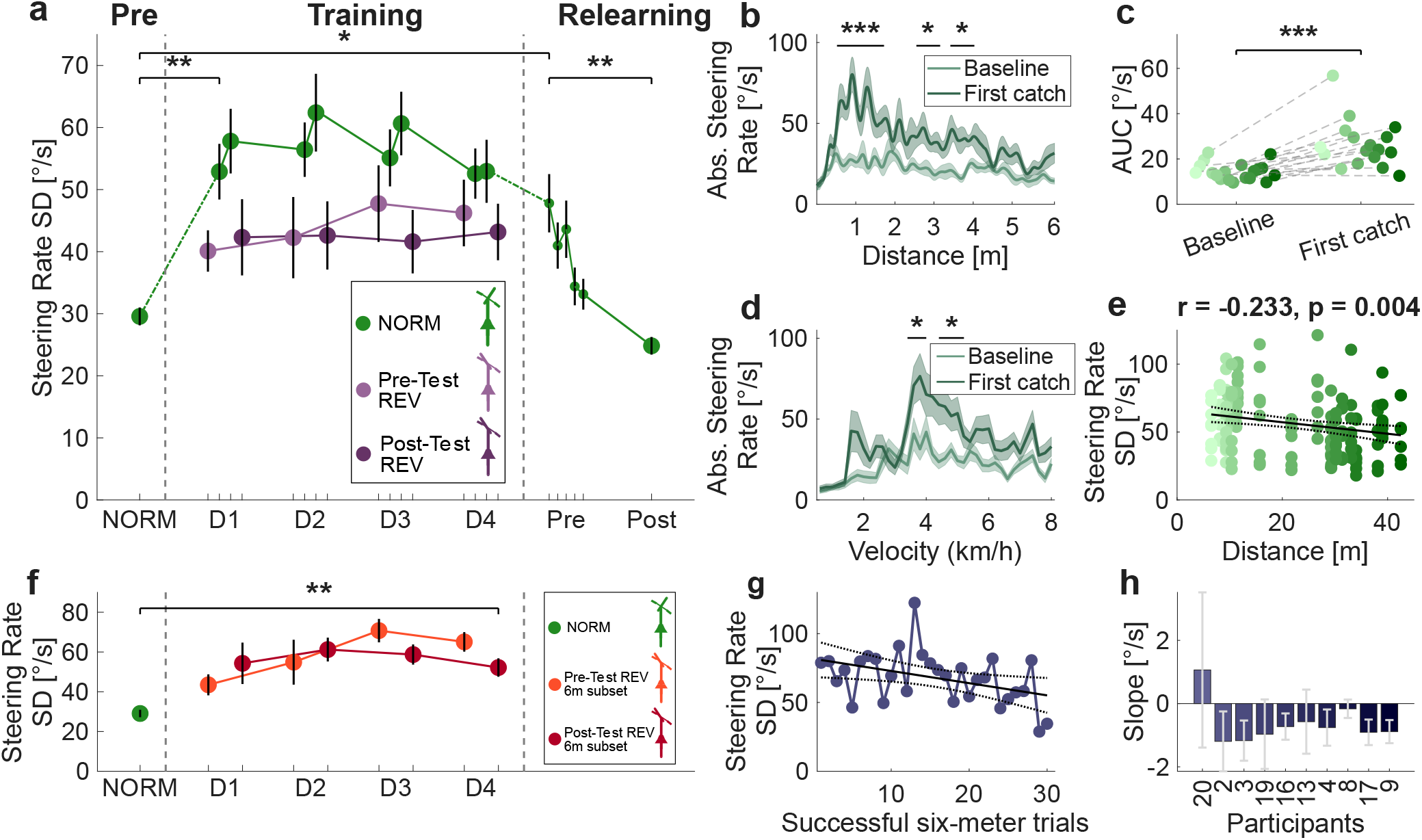
Steering rate variability on the normal and reversed bicycles demonstrates interference between the two tasks. (a) Steering rate variability (standard deviation of the steering rate) across the experiment for normal bicycle trials (green) and reversed bicycle pre-tests (light purple), and reversed bicycle post-tests (dark purple). Dotted lines distinguish experimental phases. Asterisks denote significant differences from pairwise comparisons. (b) Absolute steering rate as a function of distance for baseline trials (light green) and the first catch trial (dark green) on the normal bicycle. Black bars and asterisks indicate statistically significant clusters. (c) Area under the curve of absolute steering rate over distance for each participant in baseline (left) and first catch trials (right), with individual participants indicated by color (see Fig. 2b). (d) Absolute steering rate as a function of velocity for baseline (light green) and first catch trials (dark green). Black bars and asterisks indicate statistically significant clusters. (e) Relationship between steering rate variability in catch trials and summed reversed bicycle distance. Solid line shows linear fit, dotted lines indicate the 95% confidence interval. (f) Steering rate variability in baseline normal bicycle trials and reversed bicycle trials for the six meter subset group (n=10). (g) Linear fit of steering rate variability as a function of trials exceeding 6 m for a representative participant (participant number 9). Solid line shows linear fit, dotted lines indicate the 95% confidence interval. (h) Slopes of the linear fits for participants that achieved at least five trials exceeding 6 m, ordered and color coded by summed test distance.

Steering rate variability showed a clear measure of interference on normal bicycle riding. An ANOVA of the steering rate variability with the normal bicycle yielded a significant main effect of time (F_1.798,28.761_=12.127, p<0.001, 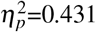). In particular, we observed a significant increase from the baseline normal bicycle trials to the first catch trial in the steering rate variability (Baseline: 30.37 ±1.51 °/s, First Catch: 53.57 ±5.11 °/s, t_16_=−4.718, p=0.001, Cohen’s d=−1.480, Fig. 3a). This indicates substantial interference with normal bicycle riding already after five minutes of reversed bicycle practice, which corresponds to an average of 90.55 ±5.83 meters across participants. Interestingly, this steering rate variability remained elevated across subsequent catch trials and only decreased significantly over the relearning phase with the normal bicycle (t_16_=4.056, p=0.006, Cohen’s d=1.480).

The steering rate variability indicates substantial interference with normal bicycle riding in these catch trials, but it does not show at which time point during a trial the interference occurs. To investigate this, we compared the absolute steering rate between the final baseline trial and the first catch trial over both the distance of the trial (Fig. 3b) and the forward velocity (Fig. 3d). Cluster-based permutation tests revealed significant increases in absolute steering rate at multiple phases of the trial and across velocity ranges during the first catch trial compared to baseline. Furthermore, we compared the area under the curve in the absolute steering rate over distance and found a significant difference between the baseline trial and the first catch trial (t_16_=−5.622, p<0.001, Cohen’s d=−1.364, Fig. 3c). While the overall absolute steering rate was generally higher across the whole distance, it can be seen that the largest difference occurred at the beginning of the trial after the push-off phase (Fig. 3b).

To assess if interference depends on the amount of achieved distance with the reversed bicycle, we then correlated the steering rate variability in catch trials to the sum of test distances (pre and post) with the reversed bicycle. We observed a small negative correlation (r=−0.233, p=0.004, Fig. 3e) indicating that greater distance achieved with the reversed bicycle might be associated with slightly less interference with normal bicycle riding.

While steering rate variability did not appear to change with the reversed bicycle over the course of the experiment (Fig. 3a, purple), a slightly different pattern can be observed when focusing on the subset of participants who eventually achieved at least five trials for the full six meters on the reversed bicycle (six meter subset group, n=10). In this subset, steering rate variability looked to initially increase, followed by a slight decrease towards the end of the experiment. (Fig. 3f). To assess whether steering rate variability decreased between the first and final six-meter trials in these participants, we performed a linear regression on the steering rate variability as a function of each participant’s successful 6 meter trials (Fig. 3g) to calculate their slope (Fig. 3h). Across this subset of participants, these slopes were predominantly negative, indicating that after successfully riding for six meters, the steering rate variability decreased in subsequent trials, reflecting improved control of steering. However, despite this improvement, participants still showed significantly higher steering rate variability in their final reversed bicycle trials compared to their baseline normal bicycle trials (t_9_=4.858, p=0.001, Cohen’s d=1.619, Fig. 3f) suggesting incomplete adaptation of steering control. Again, the steering rate variability in the normal bicycle trials was still further elevated, suggesting strong interference of the reversed bicycle on this long-learned skill.

### Balance-related variability measures

Next, we examined how a range of balance-related measures changed throughout the experiment across all participants as well as the 6 meter subset group. First, we investigated the bicycle roll rate variability (Fig. 4a, left), quantified as the standard deviation of the bicycle roll rate for each trial. We observed an increase in variability from baseline with the normal bicycle to the reversed bicycle. With the reversed bicycle, we found no significant effect on bicycle roll rate variability within days (pre-post: F_1,19_=2.553, p=0.127, 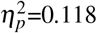) but did find a significant effect across days (days: F_1.600,30.395_=4.878, p=0.020, 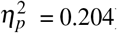). In normal bicycle trials, we observed a significant main effect of time (F_1.513,24.202_=6.552, p=0.003, 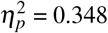), with post-hoc comparisons revealing a significant difference between the baseline trials and the first relearning trial (t_16_=−3.222, p=0.032, Cohen’s d=−1.273) and between the first relearning trial and post relearning trials (t_16_=3.845, p=0.009, Cohen’s d=1.415) indicating interference in the bicycle roll rate. To investigate if participants that were successfully able to ride the reversed bicycle (6 meter subset group; n=10) were able to produce similar roll rate variability with both bicycle types, we compared the final roll rate variability on day four post-tests to their baseline normal bicycle values (Fig. 4a, right). Although this measure appeared to decrease over the days in this subset of subjects, We still observed a significant difference compared to baseline (t_9_=−3.566, p=0.006, Cohen’s d=−1.128).

**Figure 4.**
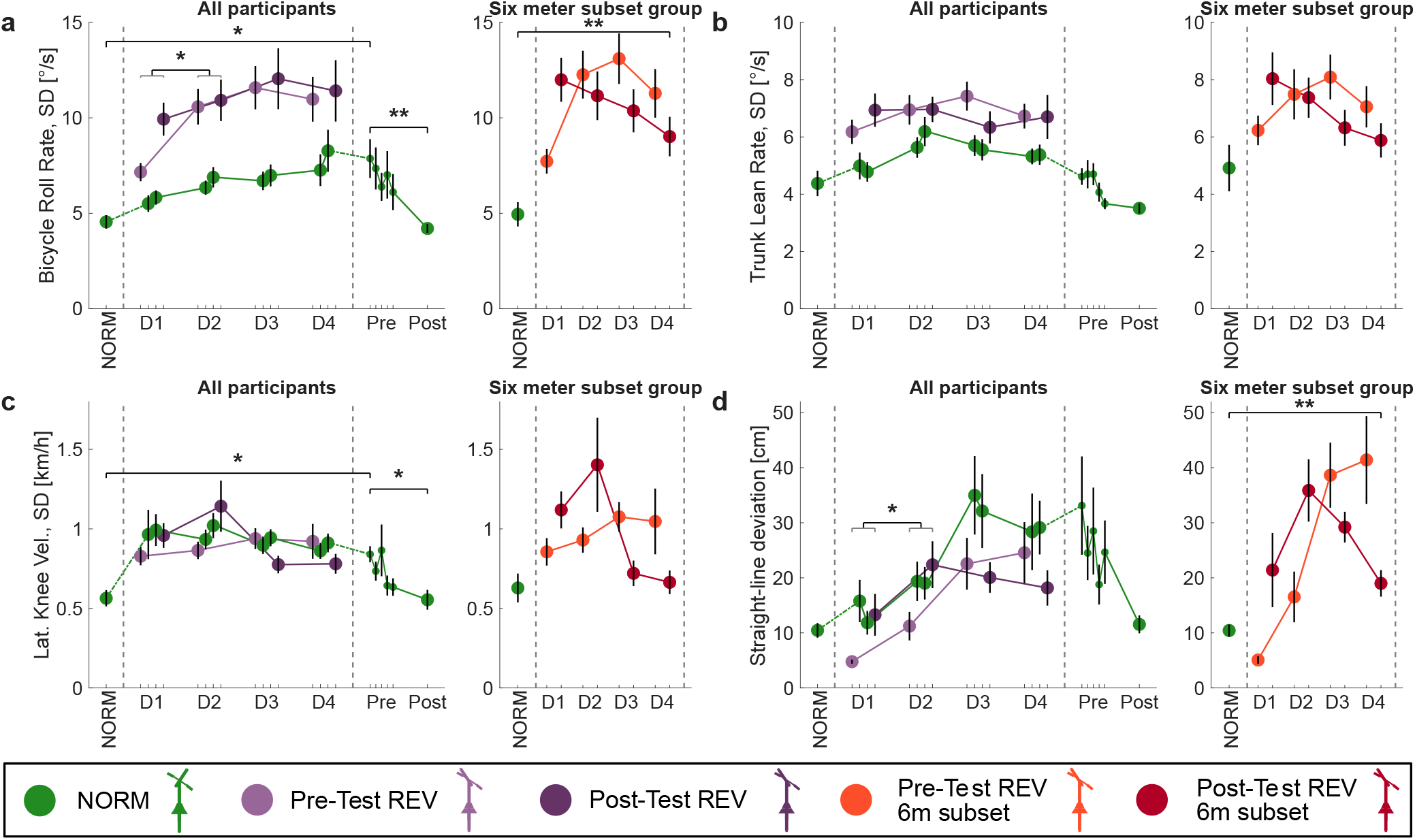
Changes in balance-related variability measures throughout the experiment for all participants as well as the six meter subset group. Trials are shown for the normal bicycle (green), reversed bicycle pre-tests (soft purple) and reversed bicycle post-tests (dark purple). In the six meter subset group, reversed bicycle pre-tests are indicated by orange, and reversed bicycle post-tests by dark red. Dotted lines distinguish experimental phases. Asterisks denote significant differences based on pairwise comparisons. When multiple pairs are significant, a gray bracket is used to group them. (a) Bicycle roll rate variability across the experiment for all participants (left) and for the six meter subset group (right, n=10). (b) Trunk lean rate variability. (c) Lateral knee velocity variability. (d) Straight-line deviation (rms of lateral deviation).

We adopted the same approach to investigate trunk lean rate variability (Fig. 4b). We observed an increase from baseline normal bicycle riding to reversed bicycle riding. However, no significant changes were found within days (F_1.726,32.800_=0.364, p=0.667, 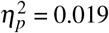) or across days (F_1,19_=0.092, p=0.765, 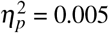) with the reversed bicycle. Furthermore, no significant differences were observed in normal bicycle riding (F_1.965,31.447_=3.088, p=0.060, 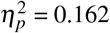). In the 6 meter subset group, we also observed no significant difference between baseline normal and post day four reversed bicycle trials (t_9_=−1.014, p=0.337, Cohen’s d=−0.321).

Lateral knee velocity variability was examined across the experiment (Fig. 4c). An ANOVA revealed no significant effect of within day (pre-post: F_1,19_=1.769, p=0.199, 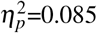) but a significant effect across days (days: F_2.219,42.157_=3.329, p=0.041, 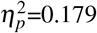). However, post-hoc comparisons revealed no significant differences between pairs (all p>0.06). In normal bicycle riding, we observed a significant main effect of time (F_1.555,24.888_=3.942, p=0.042, 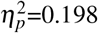). Specifically we observed a significant increase from baseline to the first relearning trial (t_16_=−3.379, p=0.023, Cohen’s d=−0.614) and a significant decrease from the first relearning trial to post relearning trials (t_16_=3.198, p=0.034, Cohen’s d=0.620). Lateral knee velocity variability was also not significantly different in normal bicycle trials and final reversed bicycle trials in the 6 meter subset group (t_9_=−0.345, p=0.738, Cohen’s d=−0.109)Fig. 4c, right).

To investigate effects on riding trajectory, we used the measure of straight-line deviation (Fig. 4d). In reversed bicycle riding, an ANOVA revealed no significant effect of within day (pre-post: F_1,19_=2.603, p=0.123,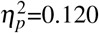) but a significant effect across days (days: F_2.066,39.251_=8.860, p<0.001, 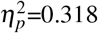) and a significant interaction (F_2.253,42.801_=3.425, p<0.037, 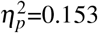). In normal bicycle riding, while the straight-line deviation appeared to increase steadily over the catch trials, an ANOVA revealed no significant main effect of time (F_1.182,18.910_=3.693, p=0.064, 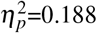). In the final reversed bicycle trials the 6 meter subset group deviated more from a straight-line compared to their baseline trials with the normal bicycle (t_9_=−3.568, p=0.006, Cohen’s d=−1.128, Fig. 4d, right) indicating limited controllability with the reversed bicycle.

On two occasions, the reversed bicycle was equipped with training wheels to support the participant against lateral falls. We compare these reversed bicycle slalom trials with training wheels to normal bicycle slalom trials and reversed bicycle slalom trials (Supplementary Fig. S2). To examine differences across the measures we fit a linear mixed model to the parameters of achieved distance, bicycle roll rate variability, bicycle roll range, and time to completion (Supplementary Note Training Wheels).

### Steering-roll coupling

Next, we examined changes in the cross correlation of steering-roll coupling as an indicator of learning during reversed bicycle riding. Previous studies^33,34,37–39^ have demonstrated a strong correlation (r=0.6-0.8) between the bicycle roll and steering adjustments used to correct that tilt, with a time delay of 90-140 ms. During normal bicycle riding, we observed a clear coupling between bicycle roll and steering (Fig. 5a). However, this relationship changes when riding the reversed bicycle successfully (Fig. 5b). In normal bicycle riding, this cross correlation was comparable to values from previous experiments (r =0.64 ±0.12, Fig. 5c). An ANOVA revealed significant differences in cross correlation in normal bicycle riding throughout the experiment (F_1.613,25.810_=4.4230, p=0.003, 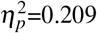) with a significant difference between baseline and the first relearning trial (t_16_=3.071, p=0.044, Cohen’s d=0.970, Fig. 5c), suggesting possible interference from the reversed bicycle. With the reversed bicycle we find a reduced cross-correlation as expected due to the reversed steering. An ANOVA on the cross correlation with the reversed bicycle yielded a main effect of within day (pre-post: F_1,19_=7.971, p=0.011, 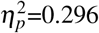) and across days (days: F_1.912,36.326_=20.262, p<0.001, 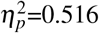). This showed that the cross-correlation for reversed bicycle riding became more negative with further practice. The final cross correlation on post-tests on day four was r = −0.46 ±0.09. However, there was high variability across participants in these final cross correlation values (Fig. 5d).

**Figure 5.**
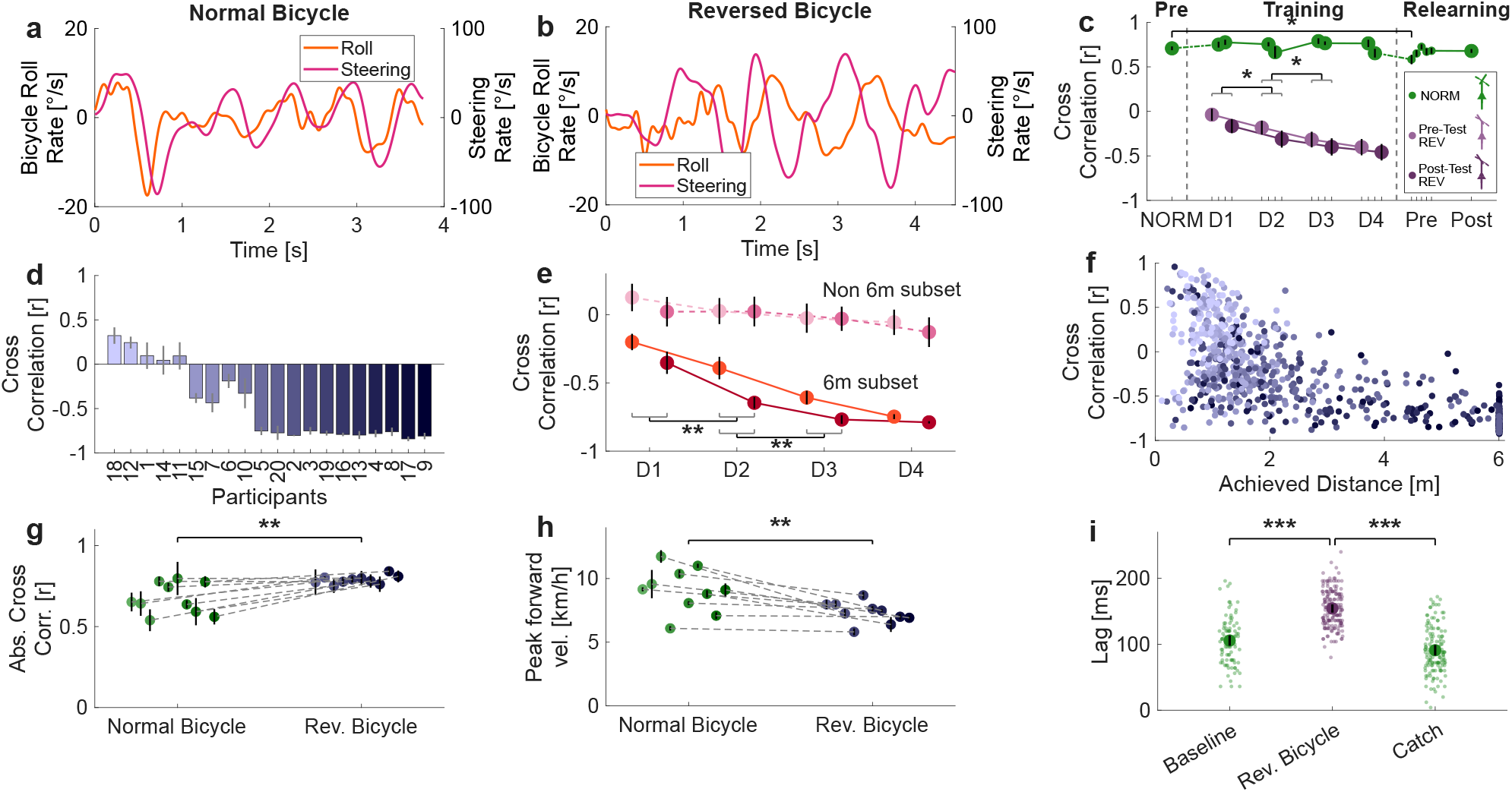
The introduction of the reversed bicycle drives changes in the cross correlation between bicycle roll rate and steering rate. (a) Example of a normal bicycle trial showing the bicycle roll rate (left y-axis; orange) and steering rate (right y-axis, pink) over time. (b) An example of a six-meter trial with the reversed bicycle. (c) Cross correlation across the experiment for normal bicycle trials (green), reversed bicycle pre-tests (light purple) and reversed bicycle post-tests (dark purple). Dotted lines distinguish experimental phases. Asterisks denote significant differences of pairwise comparisons. When multiple pairs are significant, a gray bracket is used to group them. (d) Day four post-test cross correlation (±s.e.m.) for each participant. (e) Cross correlation for the 6 meter subset group (n=10, orange and red lines) and participants outside the 6 meter subset (n=10, pink dotted lines) shown for pre-tests and post-tests. (f) Cross correlation coefficient of all reversed bicycle trials as a function of achieved distance on that trial, with colors denoting the individual participants as in Fig 2b. (g) Absolute cross correlation in baseline normal bicycle trials and day four post-test reversed bicycle trials for the six meter subset group. (h) Peak forward velocity in baseline normal bicycle and day four post-test reversed bicycle trials for the six meter subset group.(i) Time lag between roll rate and steering rate in baseline, reversed bicycle, and catch trials (mean and s.e.m.). Small dots indicate individual trials.

For the subset of participants that successfully achieved six meters for at least five trials with the reversed bicycle (six meter subset group, n=10), we observed a strong transitioning to negative steering-roll coupling with training (Fig. 5e, orange and red lines). This learning of the coupling of the reversed bicycle was confirmed with an ANOVA showing significant main effects of within day (pre-post: F_1,9_=40.378, p<0.001, 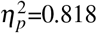) and across days (days: F_3,27_=52.589, p<0.001, 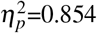). Participants outside the 6 meter subset showed cross correlation coefficients near zero throughout the experiment (Fig. 5e, pink dotted lines). More generally, across all participants we observed a clear relationship between the distance achieved in reversed bicycle riding and the cross correlation coefficients, indicating that a transition to negative cross correlation is necessary to ride the reversed bicycle successfully (Fig. 5f).

Interestingly, for participants in this six meter subset group, absolute cross correlation was significantly higher during reversed bicycle riding than during normal bicycle riding (t_10_=−3.593, p=0.005, Cohen’s d=−1.083, Fig. 5g). This might suggest that these participants are relying even more on this coupling between roll and steering to maintain the reversed bicycle upright. However, one potential contributing factor to this difference might be the variation in riding speed between conditions^33^. Indeed, we find a significant difference in the peak forward riding speed between bicycle types in this subset of participants (t_9_=3.981, p=0.003, Cohen’s d=1.259, Fig. 5h). To account for this, we fit a linear mixed-effects model with absolute cross correlation peak as the dependent variable and bicycle type and peak forward speed as fixed effects. We found significant main effects for peak forward speed (F_1,48.68_=11.756, p=0.001) and bicycle type (F_1,89.69_=7.215, p=0.009). Estimated marginal means showed that absolute cross correlation remained higher for the reversed bicycle (M=0.775, 95% CI [0.734, 0.815]) than for the normal bicycle (M=0.703, 95% CI [0.662, 0.744]) even after taking into account the differences in speed. Together, these results show that although riding speed contributes to variation in steering-roll coupling, it does not fully explain the increased absolute cross correlation observed on the reversed bicycle. Potentially, on the reversed bicycle, riders rely more strongly on a tight coupling between roll and steering while alternative control strategies, often used in normal bicycle riding, might be limited at the performance level achieved in this experiment.

We also analyzed the amount of time lag in the cross correlation between bicycle roll rate and steering rate (Fig. 5i). In normal bicycle riding we observed a time lag of 105.56 ±7.80 ms in the baseline trials and 94.57 ±8.57 ms in catch trials, whereas in trials with the reversed bicycle the lag was 156.43 ±10.23 ms. To test whether peak forward speed or achieved distance could explain this difference in the time lag in addition to bicycle type, we fit a linear mixed-effects model with time lag as the dependent variable. Bicycle type (normal bicycle baseline, reversed bicycle, and catch trials), peak forward speed, and distance were included as fixed effects. In the model, peak forward speed had no significant effect (F_1,833.69_=0.175, p=0.676), whereas distance had a significant effect (F_1,895.05_=5.316, p=0.021). In addition, bicycle type had a significant effect (F_1,907.37_=191.623, p<0.001). Specifically, the estimated marginal means showed that time lag was substantially higher for the reversed bicycle (M=160.811, 95% CI [148.905, 172.717]) compared with the normal bicycle baseline (M=103.392, 95% CI [89.301, 117.483]) and catch trials (M=92.965, 95% CI [80.024, 105.905]). Post-hoc pairwise comparisons confirmed the difference with both baseline (Δ=57.4 ms, z=12.01, p<0.001) and catch trials (Δ=67.8 ms, z=18.62, p<0.001). This shows that the differences in time lag between bicycle types are robust and cannot be explained by differences in peak speed or achieved distance.

## Discussion

Twenty participants trained to ride a reversed bicycle over four days, while interference on the normal bicycle was assessed. Around half of the participants successfully learned to ride the reversed bicycle over six meters, whereas others did not significantly increase their distance, showing the high inter-individual variability. However, even for successful participants, the balance-related performance measures with the reversed bicycle did not approach those with the normal bicycle. Nevertheless, the more successful riders showed a reorganization of the steering-roll coupling allowing them to successfully ride the reversed bicycle, although the time-lag between these measures was increased compared to the normal bicycle. There was interference on the normal bicycle riding even after only 5 minutes of training with the reversed bicycle, which remained throughout the four days, with increased steering rate variability, increased bicycle roll, increased trajectory deviation, and decreased overall riding distance.

Many studies have shown that learning one task can be generalized to similar tasks^3–6^. For example, learning to ride a bicycle might help with learning to ride a motorcycle. Within tasks, such as changing bicycle types^7^, this switching of controllers occurs quickly, supporting the idea of structural learning^3^. One might therefore expect that switching from a normal bicycle to a reversed bicycle would also be rapid. However, this is not the case. None of the participants achieved more than 1.5 meters in the first trial. Therefore, by changing one major detail of the bicycle (the inverse steering), we completely disrupt riding performance. Why is it so difficult to learn? Studies investigating mirror reversals show similar results. When the trajectory of the hand is mirrored along an axis such that the visual feedback is opposite to the actual movement, performance is disrupted similarly^41,42^. Although participants are able to adapt to this mirror-reversal, the reaction times to produce an appropriate action increase^40–44^. This increase in reaction time suggests that participants rely more on cognitive control strategies during mirror-reversal tasks, rather than rapid feedback responses. Furthermore, responses to cursor jumps under mirror-reversed feedback are still initially directed toward the incorrect direction, with delayed corrections to the appropriate direction^40,42,45^. Indeed, we have a habitual tendency to generate baseline patterns of correction when rapid adjustments are necessary^46^. In reversed bicycle riding, this would result in corrective actions appropriate for normal bicycle riding, which increases the error under the reversed steering dynamics.

With training, ten participants successfully rode the reversed bicycle for six meters at least five times during the testing (six meter subset group). This is in line with findings from Magnard and colleagues, where eleven out of twenty participants rode twenty meters at least once^13^. In both studies, many participants did not manage to ride even once successfully over the time course of the experiment. Similar behavior has been found when reaching under inverting prisms^47^, where the authors distinguished two groups of participants. Even in this less complex motor task, some participants adapted to the inversion and reduced reaching error while others did not improve or even got increasingly worse^47^. Furthermore, other studies on mirror reversal found higher inter-individual variability^41,44^ compared to adaptation studies^41^.

Despite successfully riding the reversed bicycle, participants in the six meter subset group still showed limited performance across multiple parameters compared to their baseline normal cycling. Successfully riding the reversed bicycle was characterized by increased steering rate variability, greater variability in the bicycle roll rate, and larger deviations from a straight-line trajectory. In addition, peak forward velocity was consistently lower, indicating a more cautious riding strategy. We also assessed control capability of the reversed bicycle with slalom trials on the final day, which require trajectory modulation in addition to balance control. As expected, most participants achieved less forward distance on these trials reflecting incomplete control over the trajectory. However, three participants successfully completed all five slalom trials, suggesting the emergence of goal-directed control over the riding trajectory. These observations align with previous findings that reported comparable handlebar oscillation power between both bicycle types in expert reversed bicycle riders, indicating that with extensive practice, aspects of performance could reach normal bicycle skill level^13^.

During normal bicycle riding, roll rate and steering rate are positively correlated^33,37,39^, as steering into the same direction as the bicycle tilt acts to restore balance. As reversed bicycle riding would be expected to rely on the same control strategy, successful performance could arise from a simple sign change of this coupling. If this sign could be immediately reversed in our controller, we would rapidly learn to ride the reversed bicycle. However, participants instead initially show a reduction of the cross-correlation towards zero. This steering-roll coupling then only gradually transitions to negative values with further training for some participants. Specifically, training with the reversed bicycle led to a progressive reduction in the cross correlation, and in participants who learned to ride successfully, an eventual reversal of the coupling. In contrast, the other ten participants who did not learn the reversed bicycle to this level, showed a cross-correlation near zero throughout the experiment. This might reflect the difficulty these participants had in learning to reverse this sign of the coupling.

Not only did the six meter subset group show a reversal of the steering-roll coupling, but the absolute cross correlation values during reversed bicycle trials were higher than in their normal bicycle trials. This indicates that they learned a stronger coupling between steering and roll dynamics. Indeed, there are lower cross correlation values in expert riders on training rollers compared to novice riders^33^, which could indicate greater control flexibility through contributions of additional strategies (e.g. trunk lean), resulting in reduced reliance on steering to keep the bicycle upright. Aligning with this, the higher absolute cross correlation values observed during reversed bicycle riding in our study may reflect this reduced control flexibility. This potentially indicates a greater reliance on steering for balance, suggesting that they may have not yet refined their control over trunk lean or knee motion with the reversed bicycle.

We found a consistent difference in time lag between the two bicycle types, which potentially indicates a change in the control strategy of the rider rather than simple kinematic differences. Specifically, the increased time lag in reversed bicycle riding may reveal a shift in the control mechanisms perhaps resulting from a limited predictive control strategy as a result from the steering inversion. Such an increase in the lag might indicate a shift to slower voluntary corrections instead of relying on rapid reflexes or involuntary feedback responses. We assume that riders could exhibit a similar time lag with both bicycle types after sufficient training.

In normal bicycle riding there is a higher time lag between roll and steering angle in elderly participants compared to young adults (mean difference ~45 ms)^34^ (although see^39^). Additionally, in younger adults, perturbation trials led to a significant increase in time lag (mean difference ~55 ms), which could indicate a potential safety strategy to stabilize the bicycle. The increased time lag we find during reversed bicycle riding is consistent with the idea that challenging or unfamiliar control conditions result in increased time lag between steering and roll coupling, potentially reflecting a shift towards a safety-oriented strategy and greater reliance on voluntary control.

Following the pre-tests and five minutes of practice with the reversed bicycle (average distance ridden 90.17 ±6.79 meters), the first catch trial on the normal bicycle already revealed clear interference effects. Two participants were unable to maintain balance and stepped down, and nearly all participants demonstrated increased steering rate variability and lateral knee velocity variability. While steering rate variability and knee movement variability remained consistently elevated across all catch trials, later catch trials were additionally characterized by reduced riding distance and gradual increases in roll rate variability and trajectory deviation, suggesting a progressive degradation of overall balance control. The early increase in steering rate variability in the absence of strong impairments in other balance-related measures, could indicate interference between competing steering mappings. In contrast, the broader performance decline observed in later catch trials indicates this interference extends beyond steering and affects global riding stability.

Learning the reversed bicycle shows clear interference on normal bicycle riding which increases as the training of the reversed bicycle progresses. Such effects are particularly found in dual adaptation tasks, such as learning two opposing force fields, where the learned dynamics mirror one another while performing a similar task^15,16^. Learning each of the tasks interferes with learning the other gives rise to anterograde and retrograde interference^16,48^. Despite such effects, it has been shown that combining the learning with certain additional contextual cues allows learning of both opposing dynamics simultaneously^18,19,49–51^. That is, with sufficient information about the context and experience, the sensorimotor system can separate the two dynamics and learn both simultaneously^52^. There is already some evidence that such dual adaptation is possible with the normal and reversed bicycle^13^. However, at the end of the four days of our study, interference between the two bicycle types is still present, despite the explicit knowledge of the participants about which bicycle they are about to ride on each upcoming trial. Indeed this matches force field studies which have shown learning of opposing force fields arises almost entirely from implicit adaptation^53^ suggesting limited effects of explicit knowledge on learning opposing dynamics.

## Conclusion

In this study, altering the steering mapping of a bicycle revealed the challenges of modifying long established bicycle control strategies. Our results suggest that successful reversed bicycle riding requires a gradual reorganization of the coupling between bicycle roll and steering rather than a simple inversion of the normal control strategy. The increased time lag observed during reversed bicycle trials may reflect a shift from predictive control toward a more voluntary control strategy in the unfamiliar mapping, suggesting that such control cannot be adapted immediately and likely requires extended practice.

Notably, even brief practice with the reversed bicycle resulted in substantial interference effects during normal bicycle trials, highlighting the short-term plasticity in a highly automated whole-body motor skill that is typically considered robust and resistant to change. This work extends principles from laboratory-based tasks to a complex, continuous motor task, demonstrating that the reversed bicycle not only challenges learning but can also rapidly interfere with the long established motor skill of riding a regular bicycle.

## Methods

### Participants

An a priori sample size calculation was conducted using G*Power^54^ based on results by Magnard et al. (2024)^13^, which suggested a medium effect size (Cohen’s d = 0.5). Assuming an *α* level of 0.05 and a desired power of 0.8, the analysis suggested a minimum required sample size of 16 participants. To account for potential loss-to-follow up, we aimed to recruit a total of twenty participants.

Twenty healthy adults (13 males and seven females, 25.8 ± 2.9 years, 74.2 ± 15.5 kg, 178.1 ± 10.9 cm; mean ± standard deviation) volunteered to take part in this experiment. All participants came to the laboratory for four visits on separate days (in a maximum time frame of ten days). Inclusion criteria were: no known neurological disorders, age between 18 and 40 years, no known disorders affecting balance or frailty, no uncontrolled weight loss (> 5% body weight over last year). In addition, participants were not allowed to ride any bicycle in their free time between the visits to ensure safety and avoid experimental interference.

The study was approved by the Ethics Committee of the Medical Faculty of the Technical University of Munich. All procedures were performed in accordance with the approved guidelines, including the Declaration of Helsinki. All participants provided written informed consent before participating in the study.

### Experimental setup

Two identical city bicycles (Corratec Soft Bow STX, 26” wheel size, Fig. 1b) were used in this experiment, with the only difference being the two gears mounted to the handlebar of the reversed bicycle to reverse the steering direction.

The experiment was performed in a gym hall equipped with a 3D motion capture setup (Vicon Motion Systems Ltd., Oxford, UK) using 16 Vicon cameras (Fig. 1a), with a sampling rate of 250 Hz. The calibration error was less than 0.5 mm. The capture volume spanned at least a length of 7 m and a width of 6 m.

Both bicycles were equipped with seven reflective markers, with three markers were placed on the handlebar and four markers on the frame (Fig. 1b). Seven additional markers were placed on the participant. One marker on the top of each foot, one marker on the outside of each knee pad approximately aligned with the rotation axes of the knee, and three markers along the spine reaching from the scapular region to the low-thoracic spine.

Participants performed different types of trials throughout the experiment: general trials, slalom trials, and free training. The general trials were performed in a straight line from the start position. For these trials the instruction was given: “Ride in a straight line until your rear wheel passes the marked area”. The marked area was 7.5 m away from the contact point of the front wheel. In between the start and finish position, tape spots on the floor indicated the straight line. In post-processing, the trials were cut to a maximum of 6.0 m, but we instructed participants to ride more distance to avoid breaking before passing the six meters. If participants stepped down the trial was finished. Most testing trials throughout the experiment were these ‘general trials’, unless otherwise specified.

On occasional trials, participants were asked to ride in a slalom. For these trials, two cones were placed on the straight line at distance 2.25 m and 4.5 m away from the front wheel in the start position, respectively. Participants were asked to always start going to the left, cross the cones from left to right, and then ride back to the marked straight line. Again, if participants stepped down the trial was considered to be finished at that point. The trial was also considered to be finished if the cone was moved or if participants passed the cone on the wrong side.

On two occasions, participants performed the slalom task using the reversed bicycle with training wheels. These wheels intended to support the balance of the bicycle to lateral falls. We note that the training wheels were unable to fully support the combined mass of the bicycle and participant, therefore we asked participants to step down to prevent equipment damage.

In free training sessions participants were given time to practice riding the reversed bicycle. Participants were asked to stay inside the marked capture volume (Fig. 1a). During this period, we continuously assessed the distance ridden by tracking one of the frame markers.

### Experimental procedures

After providing written informed consent, participants were asked to fill out a short questionnaire on physical activity and bicycling exposure. The first question was on physical activity “How many hours per week do you engage in physical activity?” with the answer options: “Less than one hour”, “one to three hours”, “three to five hours”, and “more than five hours”. The second question asked about riding frequency: “How often did you use the bike in the past 12 months (e.g., commute to work, shopping)” with the answer options: “Daily”, “Several times per week”, “Once a week”, “Less”, and “Never”. The third question asked about the ridden distance: “How many kilometers did you ride the bike on average per week in the past 12 months?” with the answer options: “Less than 10 km”, “10 - 30 km”, “30 - 50 km”, “50 - 100 km”, and “More than 100 km”. After filling out the questionnaire, participants were asked to wear safety gear including a helmet, elbow-, and knee pads. Markers were then placed on the participants. Participants familiarized themselves with the normal bicycle by riding freely until they felt comfortable with the bicycle. The saddle height was adjusted to the participant’s comfort.

On day one, participants started with the normal bicycle. We first captured five general trials as a baseline, followed by five slalom trials (all with the normal bicycle). Next, five pre-tests were performed using the reversed bicycle. Participants were then given four five-minute training sessions with the reversed bicycle. After the first and second session, one normal bicycle trial (catch trial) was performed to assess interference with normal bicycle riding. We note that this trial is not a catch trial per se, as participants are aware of the switch to the normal bicycle. After the third free training session, one slalom trial using the reversed bicycle with training wheels was performed. After all free training sessions, five post-tests with the reversed bicycle were performed (Fig. 1c).

On both day two and day three, participants started with five pre-tests using the reversed bicycle followed by one catch trial with the normal bicycle. Then, two ten-minute free training sessions were given. In between the two sessions, another catch trial with the normal bicycle was performed. Following training, five post-tests were performed with the reversed bicycle.

The beginning of day four was identical to the procedure on day two and three: pre-tests with the reversed bicycle, catch trial, free training, catch trial, free training and post-tests. After the post-tests with the reversed bicycle, participants attempted five slalom trials with the reversed bicycle. Then, another slalom trial with the training wheels was performed. Afterwards, five normal bicycle trials were performed to assess early relearning, where the first of these five is similar to the other catch trials. To confirm participants can safely ride the normal bicycle, participants were asked to ride ten minutes with the normal bicycle inside the capture area. Finally, five post-tests were performed with the normal bicycle.

## Data Analysis

The trajectories of the markers were reconstructed using Vicon Nexus v.2.16 (Vicon, Oxford, UK). Further processing was done using custom-written MATLAB scripts (MATLAB R2025a). Marker positions were low-pass filtered at 10 Hz with a fifth-order, zero phase-lag Butterworth filter. Small remaining gaps (< 500 ms) were filled using linear interpolation. A total amount of 44 trials (2.9%) were excluded from further analysis beyond the distance achieved due to missing marker information.

We calculated the achieved distance by measuring forward movement of a rear frame marker. Trials were cut to a maximum distance of six-meter forward riding to ensure comparability and complete marker tracking. The start of a trial was calculated as movement velocity exceeding 0.15 m/s. The end of the trial was either passing the six meters, the event of stepping down with a foot, or riding outside of the capture volume to the right or left side (handlebar midpoint marker deviating over 2.5m from the straight line).

The steering angle was calculated using the 3D coordinates of four markers placed on the bicycle. We calculated the angle using the dot product between the handlebar orientation vector (spanning left and right handlebar) and the bicycle’s longitudinal axis (spanning from the handlebar midpoint marker to a marker on the rear bicycle frame. The steering angle was then low-pass filtered at 6 Hz with a fourth order, zero phase-lag Butterworth filter. To measure the control participants have over the steering, we calculated a measure of steering rate variability as the standard deviation of the steering rate over a trial^33,37^.

To calculate the bicycle roll angle, we utilized the handlebar midpoint marker and the rear bicycle frame marker as the bicycle’s longitudinal axis (y-axis). The x-axis was defined as the vector spanning the left and right rear bicycle frame markers. The z-axis (representing the bicycle roll axis) was calculated via the cross product of these two normalized vectors. The roll angle was calculated as the angle between this z-axis and the gravitational vector. The bicycle roll angle was then low-pass filtered at 6 Hz with a fourth order, zero phase-lag Butterworth filter.

Previous research aimed to determine the coupling between steering and bicycle roll by calculating the cross-correlation between the two variables^33,37–39^. It has been suggested that this coupling must be learned to ride a bicycle successfully^37^. To assess the development of cross-correlation in reversed bicycle riding and possible effects on normal bicycle riding, we assessed the cross-correlation by using the xcorr() function in the MATLAB signal processing toolbox. We restrained the cross-correlation peaks to be found between 20 and 300 ms lag between steering and bike roll. If no clear peak was found (13.4% of trials), the cross-correlation at lag 100 ms was taken, as this value was found to be a common lag between steering and bicycle roll^38^.

To calculate trunk lean, the marker on top of the spine was projected vertically onto the ground plane. This point was then projected onto the bicycle’s roll plane. The lean angle was calculated as the angle between the line from this projected point to the marker and the line from the same point to the ground projection of the marker. To capture the trunk lean angle relative to the bicycle, the bicycle roll angle was subtracted.

We further examined lateral knee movement by calculating the lateral distance of each knee marker to the bicycle roll plane.

We aimed to assess how much participants deviate from riding in a straight line^35^. To calculate the straight-line deviation, the handlebar midpoint marker was projected vertically onto the ground plane by the plane of the bicycle roll. Therefore, this projected marker did not move laterally if the bicycle rolls in a stationary position. Straight-line deviation was then quantified as the root mean square of the lateral deviation of this projected marker.

### Statistical analyses

Statistics were performed using JASP^55^. Statistical significance was considered at p < 0.05. Normal distribution was assessed using the Shapiro-Wilk test. We performed paired t-tests and repeated-measures ANOVAs on the data. In case the assumption of normality was violated for paired comparisons, we used Wilcoxon signed-rank tests instead. If main effects were significant in an ANOVA, post-hoc tests were performed and Bonferroni corrected. Greenhouse-Geisser correction was used if data violated the assumption of sphericity. Partial eta squared 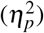 was reported as effect size for ANOVAs and Cohen’s d was reported as effect size for t-tests.

For reversed bicycle pre- and post-tests we performed a two-way (2×4) repeated-measures ANOVA with factor pre-post (2 levels: pre-test and post-test) and factor day (4 levels: day one, day two, day three, day four). To assess changes in normal bicycle performance over the course of the experiment we performed a one-way repeated-measures ANOVA with four levels (1. Baseline normal bicycle trials, 2. First catch trial, 3. First relearning trial, 4. Post ten minute relearning training trials).

Linear mixed-effects models were fit where appropriate. Models included participant as a random intercept. Fixed effects depended on the specific analysis. Degrees of freedom were estimated using the Satterthwaite approximation. Post-hoc pairwise comparisons were performed on estimated marginal means and corrected for multiple comparisons where applicable.

To quantify differences between the steering rate in baseline trials and the first catch trial, we present the absolute steering rate over distance (Fig. 3b) by interpolating the trials to 1000 data points according to distance. To statistically compare, we first calculated the area under the curves for each participant and performed a paired t-test. Furthermore, we performed a cluster-based permutation test on the data^56^. Paired t-tests were computed for each data point. Neighboring significant bins were grouped into clusters, and t values were summed up across bins. We performed 10000 permutations by random assignment of condition labels to generate the null distribution. Clusters exceeding the 95th percentile of the null distribution were considered significant.

In addition, we examined how the steering rate varies over velocity between baseline trials and the first catch trial. First, velocity data was binned from 0.5 km/h to 8.1 km/h in 0.2 km/h steps. For each participant, the steering rate was averaged per bin in both the baseline and catch trial. To statistically compare differences between the trials, we conducted another cluster-based permutation test. Paired t-tests were computed per bin, with the significance threshold adapted to the degrees of freedom (because not every participant had data in each velocity bin). Neighboring significant bins were grouped into clusters, and cluster-level statistics were calculated. Again, we performed 10000 permutations to generate the null distribution and clusters exceeding the 95th percentile of the null distribution were considered significant.

Only a subset of participants managed to ride the reversed bicycle successfully for the full six meters. To investigate differences between six-meter normal bicycle and six-meter reversed bicycle trials, we used a subset of participants (six meter subset group) consisting of the participants that managed to ride six-meters successfully for at least five trials (n=10). On this subset, we performed t-tests between their baseline normal bicycle trials and their post day four reversed bicycle trials for different parameters.

The lowest p-value of all pairs is denoted by * (p<0.05), ** (p<0.01), and *** (p<0.001) in the figures.

## Data Availability

Data that support the findings of this study are available from the corresponding author on request.

## Acknowledgments

We thank all participants for taking part in the study. We also thank the Prevention Center at TU Munich for the use of the Vicon system.

## Author contributions

P.N. and D.W.F. designed and implemented the study. P.N. collected the data. P.N. and D.W.F. analyzed the data. P.N. wrote the original draft. P.N. and D.W.F. edited and reviewed the paper.

## Additional Information

### Supplementary Information

The online version contains supplementary material available at

