## Supplementary Information for "Learning a reversed bicycle disrupts predictive control and induces interference with the normal bicycle"

### Results of the questionnaire

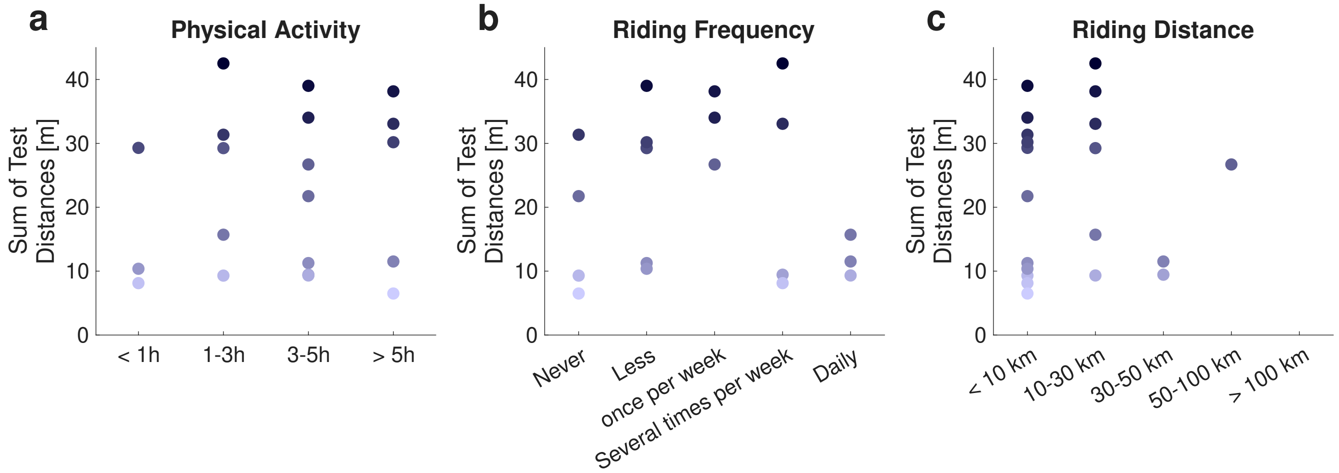


**Figure S1.** Results of the questionnaire. Each dot represents one participant. Color indicates the summed distance of all pre- and post-tests with the reversed bicycle for each participant. (a) Physical activity: Amount of hours per week. (b) Riding Frequency: Average riding frequency per week in the past 12 months. (c) Riding Distance: Amount of average distance ridden per week in the past 12 months.

### Results of the training wheel trials

After performing five trials with the normal bicycle in a straight line, participants then performed five trials with the normal bicycle in a slalom, where two cones were positioned at 2.25 and 4.5 meters from the start. On day four of the experiment, participants additionally performed five slalom trials with the reversed bicycle. In addition, to examine the effect of external stabilization, the reversed bicycle was equipped with training wheels during one trial on day one and on day four. We hypothesized that training wheels would facilitate successful performance on the reversed bicycle, but at the cost of constraining bicycle roll and thereby impairing slalom performance.

To assess performance across slalom conditions, we analyzed the distance achieved using a linear mixed model with fixed factor of test (four levels: normal bicycle slalom, reversed bicycle with training wheels day one, reversed bicycle with training wheels day four, reversed bicycle slalom without training wheels). We observed a significant main effect of test ($F_{3,57}$=19.640, p<0.001). Compared to normal bicycle trials, participants achieved shorter distances when using the reversed bicycle with training wheels on day one (Δ=-2.933 m, z=-6.426, p<0.001, Fig. S2a). Performance improved from day one to day four in the training wheel condition (Δ=2.024 m, z=4.434, p<0.001, Fig. S2a), but distance remained reduced when riding the reversed bicycle without training wheels (Δ=-1.845 m, z=-4.043, p<0.001, Fig. S2a). Notably, despite this improvement, most participants did not reach the full six meter distance in the training wheel conditions. This likely reflects limitations of the training wheel setup, as participants were instructed to step down before excessive tilting occurred, effectively constraining achievable distance. Therefore, although training wheels provided partial support, they did not fully compensate for the altered control demands of the reversed bicycle.

To assess whether training wheels constrained control of bicycle roll, we next examined bicycle roll rate variability. A linear mixed model yielded a significant main effect of test ($F_{3,51.70}$=24.009, p<0.001). Roll rate variability was lower in both training wheel conditions compared to normal bicycle trials (NORM - TW Tr. 1, Δ=-5.129 °/s, z=-6.603, p<0.001, NORM - TW Tr. 2, Δ=-4.150 °/s, z=-5.342, p<0.001, Fig. S2b). In contrast, variability increased when riding the reversed bicycle without training wheels relative to the day four training wheel condition (Δ=4.150 °/s, z=5.342, p<0.001), suggesting a less constrained and more actively controlled use of roll.

Because roll variability may be influenced by failed trials, we further quantified the range of bicycle roll in successful trials only (i.e., trials exceeding six meters). The linear mixed model again revealed a significant main effect of test ($F_{3,31.12}$=8.496, p<0.001). Roll range was reduced in both training wheel conditions compared to normal bicycle trials (NORM - TW Tr. 1, Δ=-9.552 °, z=-4.176, p<0.001, NORM - TW Tr. 2, Δ=-5.194 °, z=-3.154, p=0.010, Fig. S2c). In addition, roll range was greater in reversed bicycle trials without training wheels compared to the day four training wheel condition (Δ=5.768 ° z=2.785, p=0.032). These findings indicate that training wheels substantially restricted the available range of roll motion.

Finally, to assess how these constraints affected task execution, we analyzed the time required to reach six meters in successful trials (Fig. S2d). A linear mixed model revealed a significant main effect of test ($F_{3,31.31}$=30.611, p<0.001). Completion times were longer in the day four training wheel condition compared to both normal bicycle trials (Δ=7.127 s, z=5.650, p<0.001) and reversed bicycle trials without training wheels (Δ=5.703 s, z=3.562, p=0.002).

Together, these results demonstrate that training wheels constrained bicycle roll, reducing both roll rate variability and roll range. This restriction was associated with impaired slalom performance. We suggest that effective control of bicycle roll is critical for successful navigation in the slalom course and was hindered by the external stabilization of the training wheels.

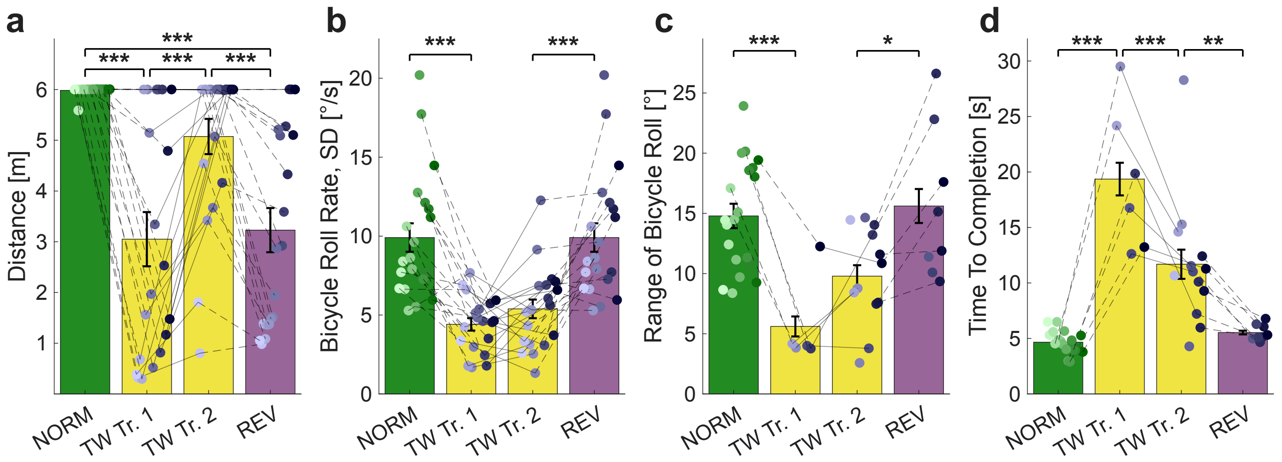


**Figure S2.** Overview of slalom trials with training wheels. NORM (far left) indicates the slalom trial on day one with the normal bicycle. TW indicates the reversed bicycle with training wheels (mid-left: day one, mid-right: day four). REV (far right) indicates the slalom trial with the reversed bicycle without training wheels on day four. The bar height indicates the mean. The black bar indicates the standard error of the mean. Individual participants indicated by color gradient. (a) Distance achieved in meters. (b) Bicycle roll rate variability as the standard deviation of bicycle roll rate. (c) Range of bicycle roll (only six-meter trials). (d) Time to Completion in seconds (only six-meter trials).
